# Effects of novelty and temporal distance on post-experience reactivation of hippocampal place cells encoding multiple environments

**DOI:** 10.1101/2023.12.17.572075

**Authors:** Taiki Yokoi, Yu Shikano, Haruya Yagishita, Yuji Ikegaya, Takuya Sasaki

## Abstract

The hippocampus plays a crucial role in consolidating episodic memories from diverse experiences that encompass spatial, temporal, and novel information. This study analyzed the spike patterns of hippocampal place cells in the CA3 and CA1 areas of rats that sequentially foraged in five rooms, including familiar and novel rooms, followed by a rest period. Across multiple rooms, the generation of place fields by CA1 place cells was coordinated with other place cells. In the subsequent rest period, CA3 place cells that encoded novel environments exhibited stronger and more coordinated reactivation during sharp wave ripples (SWRs) than CA1 place cells. In contrast, CA1 place cells that encoded more recent environments exhibited stronger SWR-associated reactivation, independent of spikes of other cells, with weaker influences from novelty compared to CA3 place cells. These results suggest that post-experience SWR-associated reactivation of CA3 and CA1 neurons primarily processes novelty-related and temporal distance-related aspects of memory, respectively.

## INTRODUCTION

From sequential experiences, animals need to extract and process information related to the temporal, spatial, and novel aspects of their experiences. The hippocampus plays a crucial role in episodic memory ^1,2^ and contains neurons known as place cells that represent spatial information by firing at particular locations in an environment ^3,4^. In different environments, spatial encoding patterns of place cells have been shown to differ between CA3 with recurrent networks and CA1 with feed-forward networks ^5,6^. In CA3, distinct sets of hippocampal place cells independently encode spaces in different environments ^5,7,8^, enabling orthogonalized spatial representations to minimize the interference of spatial maps by place cells. In CA1, place cells more frequently exhibit spatial representations within an environment, and a larger proportion of neurons consequently exhibit spatial representations across multiple environments compared with the CA3 ^5,6^. From a temporal perspective, the spatial firing patterns of CA3 neurons are more stable over time than those of CA1 neurons ^9,10^. The distinct stability of spatial encoding between the two subregions may underlie their unique roles in processing temporal information, such as temporal distance from experiences.

During rest/sleep periods, following the experience of an environment, place cell ensembles are synchronously reactivated, typically during sharp-wave ripples (SWRs) in local field potential (LFP) signals ^11-23^, which are believed to serve as neurophysiological substrates for memory consolidation. SWRs originate from the CA3 recurrent circuit, involving the reactivation of experienced information ^24^, and subsequently establish coordinated neuronal spike assemblies in the downstream CA1 area ^25,26^. Crucially, both CA3 and CA1 place cells that encode novel than familiar environments exhibit more substantial increases in firing rates and firing associations during SWRs ^20,22,27,28^, potentially offering an efficient mechanism for integrating novel information into hippocampal circuits.

The significance of hippocampal reactivation raises more intricate questions regarding how reactivation occurs when place cells encode multiple environments consisting of a combination of various spatial, temporal, and novel/familiar information. Moreover, a crucial question at the cell ensemble level is how place cell reactivation is coordinated with or interfered with by other neurons. To clarify these issues, we recorded the spike patterns of hippocampal CA3 and CA1 place cells as rats sequentially experienced five different rooms, alternating between familiar and novel rooms, and then rested in the familiar box. After confirming the spatial representation patterns across multiple rooms by CA3 and CA1 place cells, we analyzed how post-experience SWR-associated reactivation of the place cell ensembles was affected by the novelty of the environments and temporal distance from the experiences.

## RESULTS

### Experimental timeline

Rats were trained daily for one week to forage for food rewards in three rooms (room 1, 3, and 5) with different shapes, each placed in a different location surrounded by different external cues. Subsequently, these rooms were used as familiar rooms (Fig. 1A and Fig. S1). After the rats were implanted with the tetrode assembly, they were again trained daily in the three identical rooms while the tetrodes were advanced to the hippocampal CA1 pyramidal cell layer. On the day of CA1 neuron recording, the rats first rested in a familiar box (pre-rest) and successively performed random foraging in five rooms from room 1 to 5 every 10 min, each of which was flanked by a rest interval of 5 min in the same box. Finally, the rats rested again in the same box (post-rest) (Fig. 1A). Among these five rooms, room 2 and 4 were novel for the rats, situated in different settings with distinct external cues compared to the three familiar rooms (Fig. S1). After recording CA1 neuronal spikes for several days, tetrodes were advanced into the hippocampal CA3 pyramidal cell layer in the same rats. Recordings were then obtained from the CA3 neurons using the same procedures, except that room 2 and 4 were situated in completely different settings, including different cues and room coloring, so that they served as novel rooms every time.

**Figure 1.**
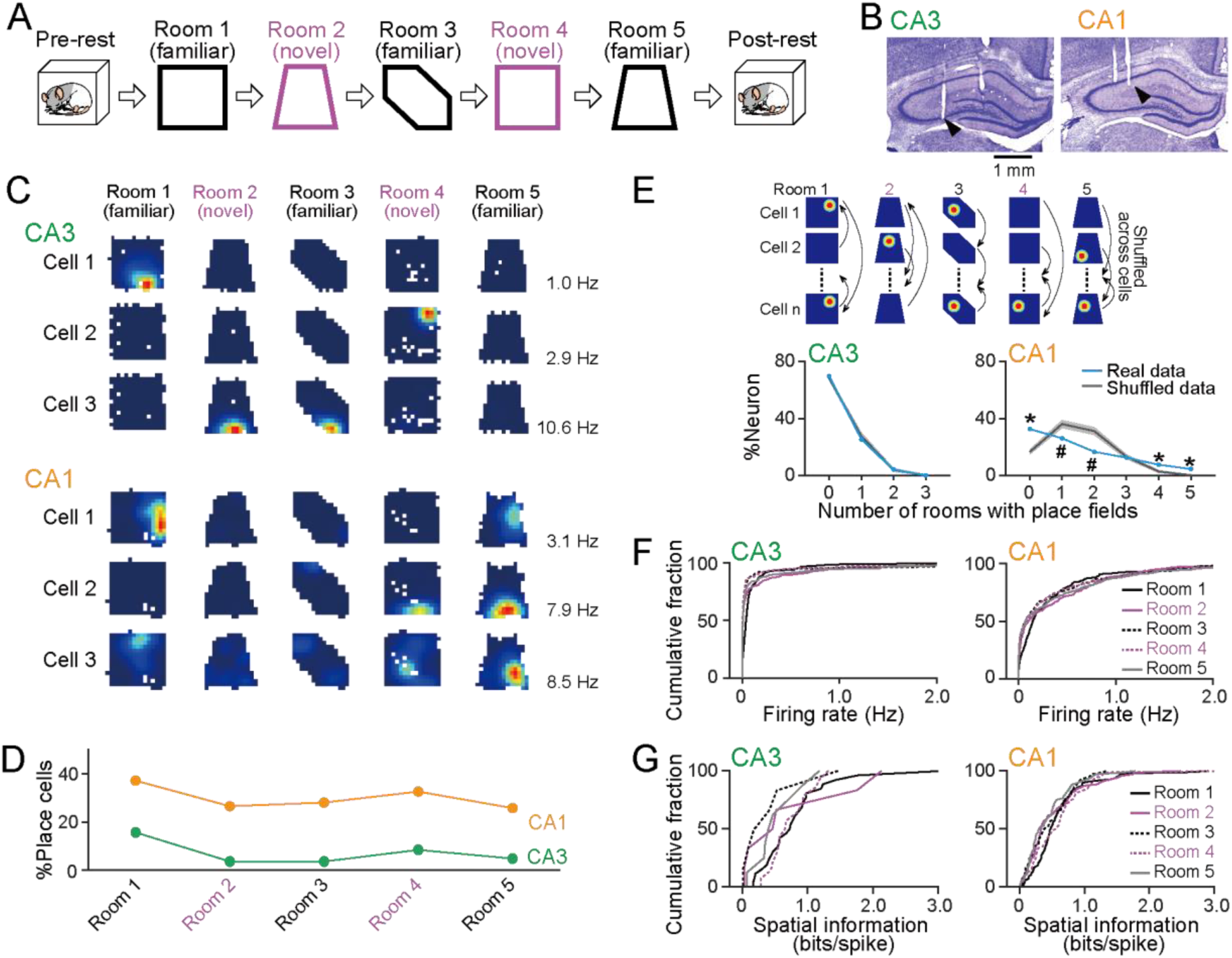
Spatial encoding of CA3 and CA1 neurons in the five rooms. (A) On the day of recording, a rat sequentially experienced three familiar, and two novel rooms. Before and after the sessions, the rat rested in a box, termed pre-rest and post-rest periods. (B) Representative images of cresyl violet-stained brain sections showing tetrode locations in the CA3 and CA1 (arrowhead). (C) Color-coded firing rate maps of representative CA3 and CA1 cells in the five rooms (blue: 0 Hz; red: peak rate indicated by the left bottom in each panel). (D) The percentages of place cells identified in each room (CA3: *n* = 166 cells; CA1: *n* = 264 cells). (E) The probability distributions of the number of rooms in which place fields were observed from single neurons. Real and shuffled data are shown in black and cyan, respectively. Data are presented as the mean ± SD. \**P* < 0.05 (real larger) and ^#^*P* < 0.05 (shuffle larger), Student’s *t*-test followed by Bonferroni correction. (F) Cumulative distributions of average firing rates in each room (CA3: *n* = 166 cells; CA1: *n* = 264 cells). (G) Cumulative distributions of spatial information of place cells identified in each room (CA3: *n* = 26, 6, 6, 14 and 8 place cells; CA1: *n* = 98, 70, 74, 86 and 68 place cells in room 1, 2, 3, 4 and 5, respectively).

### Spatial encoding by hippocampal CA3 and CA1 neurons in multiple rooms

In total, 166 and 264 putative excitatory pyramidal neurons were recorded from the CA3 and CA1 areas in 6 and 10 recordings, respectively, from five rats (Fig. 1B and Table S1). Overall, the proportion of CA1 neurons showing place fields in each room was significantly higher than that of CA3 neurons (Fig. 1C and 1D; *F*1,8 = 97.7, *P* = 9.3 × 10^−6^, one-way ANOVA). In both CA3 and CA1, no significant differences were observed in the proportions of neurons showing place fields in familiar or novel rooms (Fig. 1D; CA3: χ^2^ = 0.51, *P* = 0.47; CA1: χ^2^ = 0.036, *P* = 0.85, chi-squared test). The number of rooms in which place fields were observed from single CA1 neurons was significantly higher than that from single CA3 neurons (Fig. 1E; CA3: *n* = 50 place cells; CA1: *n* = 178 place cells, *Z* = 5.74, *P* = 4.6 × 10^−9^, Mann–Whitney U test). To evaluate whether the number of rooms with place fields observed from single-place cells could be statistically explained by chance, we created shuffled datasets in which the spatial maps of individual neurons were randomly assigned to other neurons without altering the total number of place cells in each room (Fig. 1E, top). This shuffling procedure eliminated any biases in the spatial representations across multiple rooms for specific neurons. In CA3, the probability of identifying the number of rooms with place fields from single neurons did not significantly differ between the real and 1000 surrogate datasets (Fig. 1E, left; zero place fields: *n* = 116 cells, *t115* = 1.23, *P* > 0.99; one place field: *n* = 42 cells, *t41* = –0.94, *P* > 0.99; two place fields: *n* = 7 cells, *t6* = 0.30, *P* > 0.99, Student’s *t-*test followed by Bonferroni correction). In CA1, the proportion of neurons showing no place fields in any of the five rooms was significantly higher in the real datasets than in the shuffled datasets (Fig. 1E, right; *n* = 86 cells, *t85* = 9.44, *P* = 1.5 × 10^−19^, Student’s *t*-test followed by Bonferroni correction). On the other hand, the proportions of neurons showing one or two place fields out of the five rooms were significantly lower (Fig. 1E, right; 1 place field: *n* = 69 cells, *t68* = –3.75, *P* = 0.0011; 2 place fields: *n* = 44 cells, *t43* = –5.27, *P* = 1.0 × 10^−6^, Student’s *t*-test), whereas those showing four or five place fields were significantly higher, compared with shuffled datasets (Fig. 1E, right; 4 place fields: *n* = 20 cells, *t19* = 5.31, *P* = 8.0 × 10^−7^; 5 place fields: *n* = 12 cells, *t11* = 14.34, *P* = 9.2 × 10^−42^, Student’s *t*-test followed by Bonferroni correction). These results confirm that subsets of CA1 neurons preferentially maintained no spatial selectivity or generated more frequent spatial representations across multiple environments than expected by chance. No significant differences in the average firing rates were observed among the five rooms (Fig. 1F; CA3: *n* = 166 cells; CA1: *n* = 264 cells; *P* > 0.05, Mann–Whitney U test followed by Bonferroni correction for all comparisons). Moreover, no significant differences in the spatial information of place fields were observed among the five rooms (Fig. 1G; CA3: *n* = 26, 6, 6, 14, and 8 cells; CA1: *n* = 98, 70, 74, 86, and 68 cells; *P* > 0.05, Mann–Whitney U test followed by Bonferroni correction in all comparisons). These results demonstrate that the excitability and spatial selectivity of both CA3 and CA1 place cells were not considerably affected by the novelty or order of the room that the rats experienced.

### Coordinated spatial encoding by CA1 place cells across multiple environments

Next, we investigated whether CA1 place cell ensembles cooperatively encode multiple environments. Due to the limited number of CA3 neurons showing place fields in multiple rooms (Fig. 1E), the following analyses were not applied to CA3 neurons. In each room, CA1 neurons showing place fields and no place fields were labeled as “P” and “N,” respectively, and neuron pairs were classified into four types depending on their labels (P-P, P-N, N-P, and N-N) (Fig. 2A). The joint probabilities of observing each type of neuronal pair in the two rooms were computed (Fig. 2B). To assess the significance of each probability, the probability was z-scored by computing probabilities from the corresponding 1000 surrogate datasets, in which the labels of neurons were randomly shuffled across all neurons without altering the total number of labels in each room. In some rooms, joint probabilities of observing the same neuronal pair types (e.g. P-P, P-N, N-P, and N-N) were significantly higher than the chance levels as indicated by zscores of more than 2.96 (giving *P* < 0.05, followed by Bonferroni correction), while some of joint probabilities of observing different neuronal pairs were significantly lower than the chance levels as indicated by zscores of less than –2.96. These results suggest that specific pairs of CA1 neurons exhibit a preference for either forming or not forming spatial representations. To further test the cooperativity of spatial representations by CA1 neurons, a spatial correlation was computed from the spatial maps of a P-P pair in each room (Fig. 2C), and the spatial correlations of all neuron pairs were plotted for all room pairs (Fig. 2D). Some room pairs showed significantly positive correlations between the spatial correlations of P-P pairs (room 2 vs. room 5: *n* = 72 cell pairs; *r =* 0.33*, P* = 0.049; room 4 vs. room 5: *n* = 106 cell pairs; *r =* 0.37*, P* = 9.2 × 10^−4^, Pearson’s correlation analysis followed by Bonferroni correction). To further confirm cooperative spatial representations at the neuronal ensemble level, a population vector (PV) was constructed from the firing rates of all active neurons within a 7 × 7 cm location bin in a spatial map of each room (Fig. 2E, top), and PV correlations were computed from all possible pairs of location bins (Fig. 2E, bottom). All distributions of PV correlations were significantly skewed to the positive side compared to those computed from the corresponding 1000 surrogate datasets, in which firing rates were randomly shuffled across all neurons in each location bin (Fig. 2F; the lower limit of the 95% confidence interval of *Z* values was more than 0, as defined by the Mann–Whitney U test from 1000 bootstrap samples followed by Bonferroni correction). These results demonstrate that the spatial representation patterns of CA1 place cells exhibit some overlap across different environments, suggesting that specific CA1 place cell ensembles are preferentially recruited by identical spatial maps in different environments.

**Figure 2.**
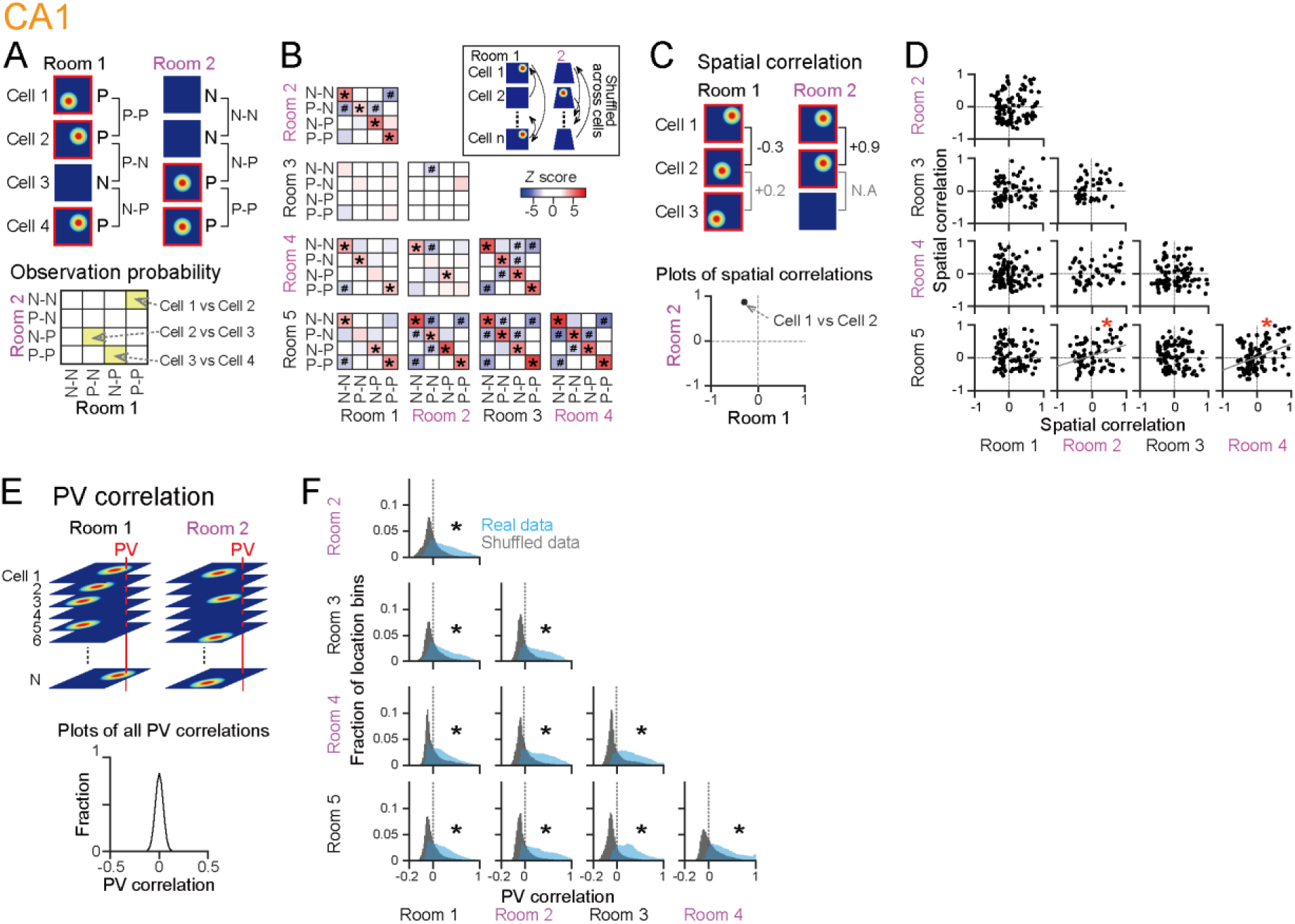
Cooperative spatial representations by CA1 neurons across multiple environments. (A) For two given rooms, joint probabilities to observe CA1 neuron pair types were computed. (B) Pseudo-colored matrices showing zscored joint probabilities from the two given rooms, constructed for all pairs of rooms. \**P* < 0.05 (positive) and ^#^*P* < 0.05 (negative), defined from zscores of more than 2.96 and less than –2.96, respectively. (C) For the two given rooms, spatial correlations of pairs of neurons were computed. (D) Spatial correlations were plotted between given two rooms, constructed for all pairs of rooms. \**P* < 0.05, Pearson’s correlation analysis followed by Bonferroni correction. (E) For the two given rooms, correlation coefficients of PVs were computed from all pairs of location bins. (F) Distributions of PV correlations between all pairs of locations in given two rooms, constructed for all pairs of rooms. The asterisk (*) represents that the lower limit of the 95% confidence interval of *Z* values was more than 0, defined by Mann–Whitney U test from 1000 bootstrap samples followed by Bonferroni correction.

### Effects of the number of spatial representations on post-experience SWR-associated reactivation of CA3 and CA1 place cells

Next, we investigated how individual hippocampal cells were reactivated in the post-rest period after the rats experienced the five rooms (Fig. 3A). Post-experience spike patterns of hippocampal cells have been shown to represent not only experience-dependent neuronal reactivation but also preconfigured neuronal population activity that emerges during pre-experience periods ^29-32^. To examine experience-dependent neuronal reactivation, we evaluated the spike patterns in the post-rest period and compared them with those in the pre-rest period. First, SWRs were detected from LFP signals during rest periods using conventional procedures ^15,19,33^. The rates of SWRs in the post-rest period were significantly higher than those in the pre-rest period (CA3; *n* = 6 recording days from 3 rats; *t5* = 5.26, *P* = 0.0033, CA1; *n* = 10 recording days from five rats; *t9* = 4.97, *P* = 7.7 × 10^−4^, paired *t* test). To compare the SWR-associated spike rates of hippocampal neurons between the pre- and post-rest periods (Fig. 3A), scatter plots were constructed separately for different numbers of rooms in which individual neurons showed place fields (Fig. 3B and 3D). CA3 neurons that had no place fields in any of the five rooms exhibited no significant differences in SWR-associated spike rates between the two rest periods (Fig. 3B, zero place fields: *n* = 105 cells, *Z* = –1.88, *P* = 0.060, Wilcoxon signed-rank test). On the other hand, CA3 place cells that had place fields in one or two rooms showed significantly higher SWR-associated spike rates during the post-rest period than those during the pre-rest period (Fig. 3B, 1 place field: *n* = 39 cells, *Z* = 3.31, *P* = 9.4 × 10^−5^; 2 place fields: *n* = 7 cells, *Z* = 2.20, *P* = 0.028, Wilcoxon signed-rank test). To quantify the strength of post-rest reactivation across different neuronal groups, we computed the SWR reactivation ratio for each cell as the ratio of the difference in the SWR-associated spike rates between the pre- and post-rest periods to the sum of these rates (Fig. 3B, bottom). Cells that generated no spikes in either the pre- or post-rest period were excluded from the analysis. The ratios ranged between –1 and +1, with higher positive and negative ratios indicating a stronger bias toward the pre-rest and post-rest periods, respectively. SWR reactivation ratios of CA3 place cells that had place fields in at least one room were significantly higher than those that had no place fields in any of the five rooms (Fig. 3C; *n* = 105 and 46 cells, *Z* = 3.37, *P* = 7.5 × 10^−4^, Mann–Whitney U test). These results confirm the robust increase in post-experience SWR-associated reactivation of CA3 place cells.

**Figure 3.**
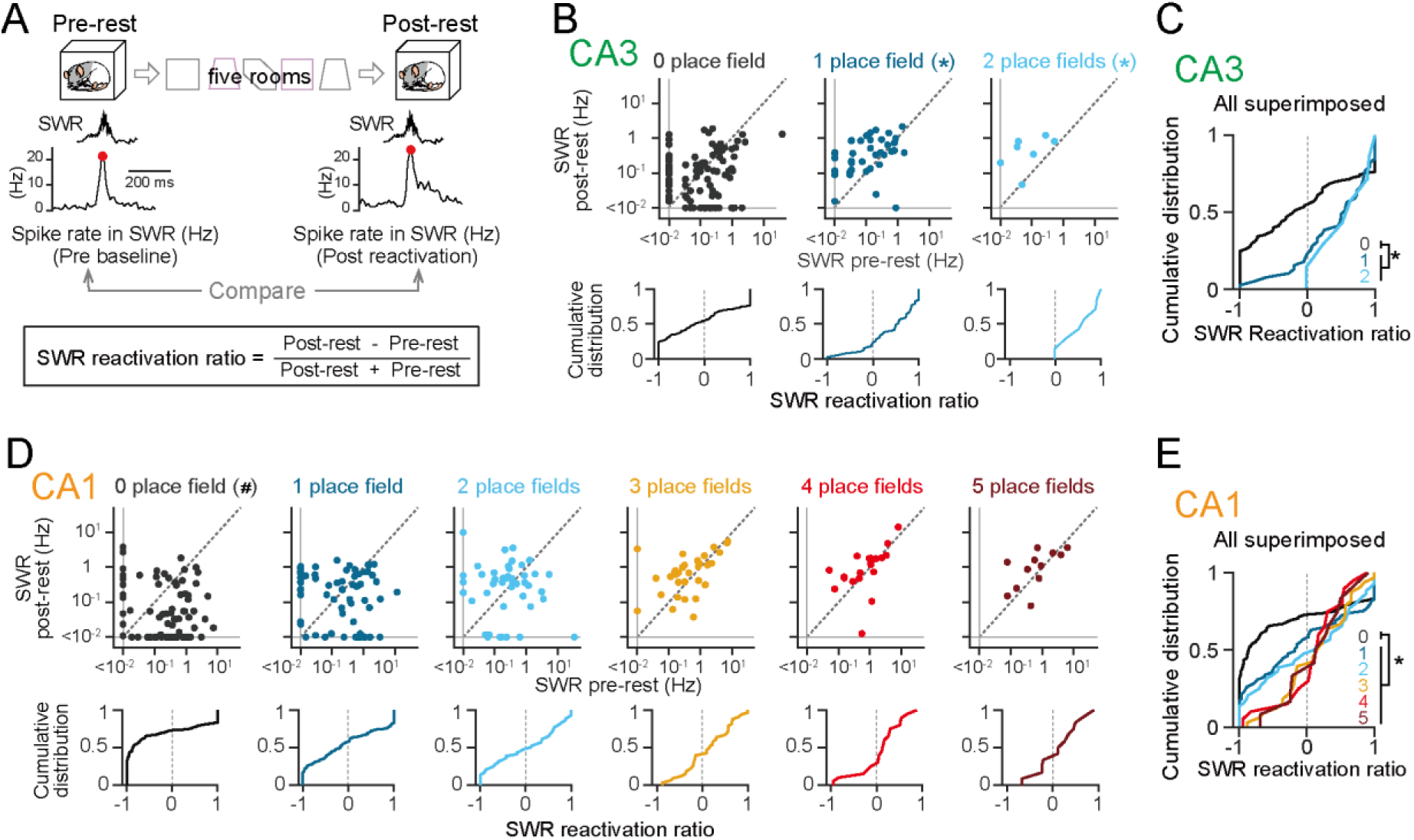
Post-experience SWR-associated reactivation of place cells showing different counts of spatial representations. (A) SWR-associated spike rates in the pre- and post-rest periods were compared. A SWR reactivation ratio was computed from each neuron. (B) (Top) Scatter plots of SWR-associated spike rates of CA3 neurons during the post-rest period against those during the pre-rest period, separately created for each number of rooms (0: *n* = 105 cells; 1: *n* = 39 cells; 2: *n* = 7 cells). \**P* < 0.05 (post larger), Wilcoxon signed–rank test. (Bottom) Cumulative distributions of SWR reactivation ratios computed from the scatter plots. (C) All distributions shown in B are superimposed. \**P* < 0.05, 0 versus 1–2, Mann– Whitney U test. (D, E) Same as B and C but for CA1 neurons (0: *n* = 85 cells; 1: *n* = 65 cells; 2: *n* = 44 cells; 3: *n* = 33 cells; 4: *n* = 20 cells; 5: *n* = 12 cells). ^#^*P* < 0.05 (pre larger), Wilcoxon signed–rank test. \**P* < 0.05, 0 versus 1–5, Mann–Whitney U test.

CA1 neurons with one or more place fields showed no significant differences between the two rest periods (Fig. 3D; one place field: *n* = 65 cells, *Z* = –1.88, *P* = 0.060; two place fields: *n* = 44 cells, *Z* = 0.55, *P* = 0.58; three place fields: *n* = 33 cells, *Z* = 0.65, *P* = 0.51; four place fields: *n* = 20 cells, *Z* = 1.42, *P* = 0.16; five place fields: *n* = 12 cells, *Z* = 0.71, *P* = 0.48,Wilcoxon signed-rank test). These results demonstrate that the number of environments in which CA1 place cells showed spatial encoding did not exert pronounced effects on subsequent post-experience SWR-associated reactivation. Importantly, CA1 neurons that had no place fields in any of the five rooms showed significantly lower SWR-associated spike rates during the post-rest period than during the pre-rest period (Fig. 3D, 0 place field: *n* = 85 cells, *Z* = –4.21, *P* = 2.6 × 10^−5^, Wilcoxon signed-rank test), and their SWR reactivation ratios were significantly lower than those of CA1 place cells that had place fields in at least one room (Fig. 3E; *n* = 85 and 174 cells, *Z* = 4.37, *P* = 1.2 × 10^−5^, Mann– Whitney U test). These results demonstrate that the SWR-associated reactivation of CA1 neurons showing no spatial representation was specifically weakened ^19^.

### Effects of the novelty and temporal distance on post-experience SWR-associated reactivation of CA3 and CA1 place cells

To test the effects of novelty and temporal distance on post-experience reactivation, the SWR reactivation ratios were analyzed separately for place cells showing place fields in each room. In CA3, place cells that had place fields in room 1, 2, 4, and 5 showed significantly higher SWR reactivation ratios than those of the other cells in each room (Fig. 4A, top; room 1: *n* = 24 cells, 95% 0.40 to 3.83; room 2: *n* = 5 cells, 0.41 to 2.07; room 3: *n* = 5 cells, –1.51 to 2.07; room 4: *n* = 12 cells, 3.06 to 3.08; room 5: *n* = 7 cells, 0.17 to 2.41, 95% confidence intervals of *Z* values defined by Wilcoxon signed-rank test from 10000 bootstrap samples). The superimposition of the cumulative distributions revealed a tendency for higher SWR reactivation ratios in the two novel rooms (Fig. 4B). These effects were further tested using a linear regression approach in which the SWR reactivation ratios of individual neurons (dependent variable) were regressed from their spatial representation patterns (predictor vectors) (Fig. 4E). For each neuron, the predictor vector consisted of binary indices indicating the presence or absence of place fields (1 or 0, respectively). Linear regression yielded the coefficient *β*, representing the strength and direction of the effect of spatial encoding in individual rooms. For CA3 place cells, room 2, 4, and 5 had higher *β* values than room 1 and 3, demonstrating that the two novel rooms and the latest rooms had stronger effects on the SWR reactivation ratios (Fig. 4F, left; *n* = 46 place cells).

**Figure 4.**
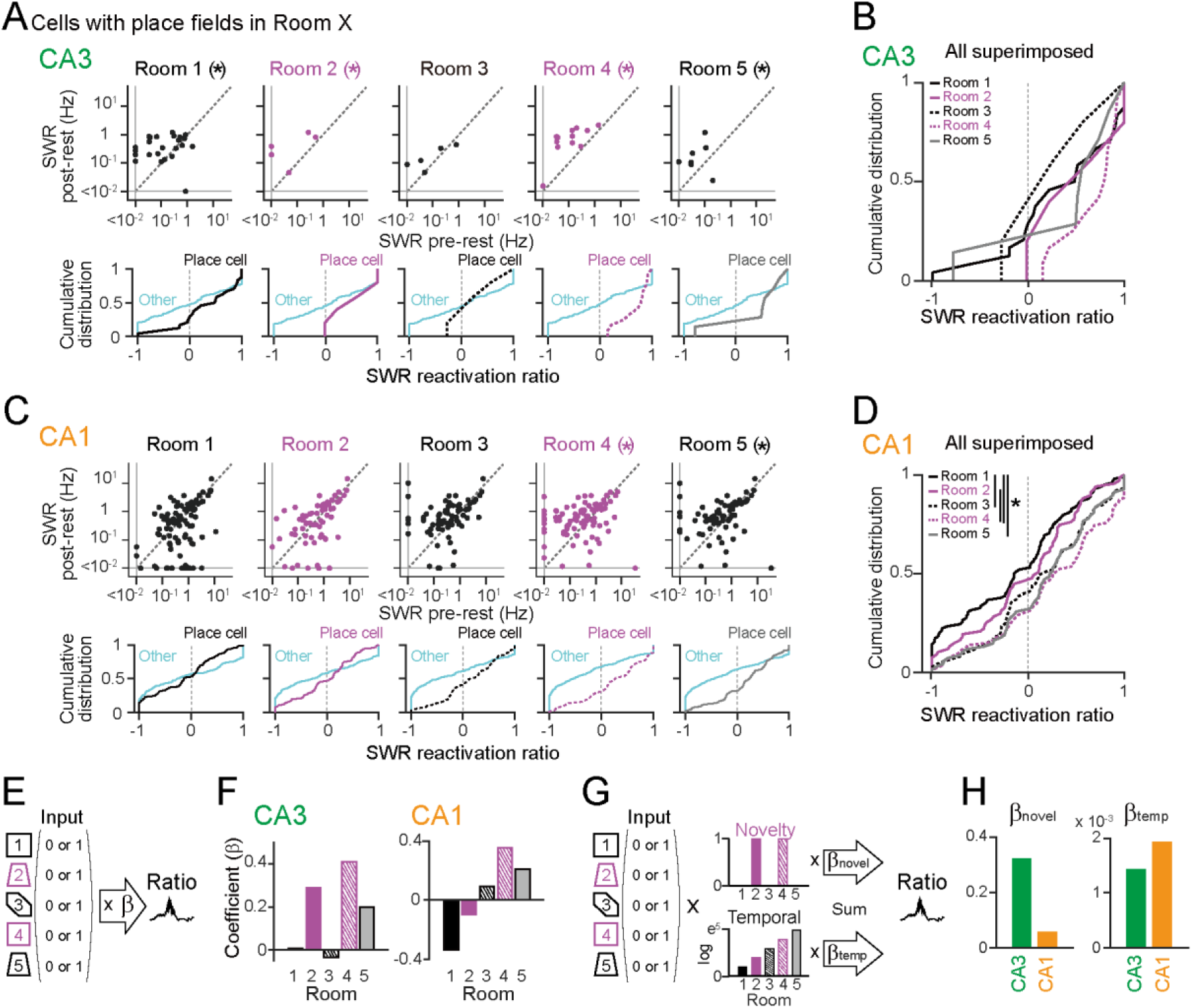
The effects of novelty and temporal distance on post-experience SWR- associated reactivation of place cells. (A) (Top) Scatter plots of SWR-associated spike rates of CA3 neurons during the post-rest period against those during the pre-rest period, separately created for neurons that had place fields in each of the five rooms (room 1: *n* = 24 cells; room 2: *n* = 5 cells; room 3: *n* = 5 cells; room 4: *n* = 12 cells; room 5: *n* = 7 cells). The asterisk (*) represents that the lower limit of the 95% confidence interval of *Z* values was more than 0 (post larger), defined by Wilcoxon signed-rank test between pre and post from 10000 bootstrap samples. (Bottom) The corresponding cumulative distributions of SWR reactivation ratios. Black or magenta lines represent neurons that had a place field in that room, while cyan lines represent the other neurons that had no place fields. (B) All distributions shown in A are superimposed. (C) Same as A but for CA1 neurons (room 1: *n* = 98 cells; room 2: *n* = 67 cells; room 3: *n* = 73 cells; room 4: *n* = 86 cells; room 5: *n* = 68 cells). (D) Same as B but for CA1 neurons. \**P* < 0.05, Mann–Whitney U test followed by Bonferroni correction. (E) The SWR reactivation ratios were linearly regressed against binary-coded spatial representation patterns. (F) Regression coefficients *β* for the individual rooms computed from CA3 and CA1 place cells (*n* = 46 and 174 cells). (G, H) Using multiple linear regression, coefficients *βnovel* and *βtemp* were computed from CA3 and CA1 place cells (*n* = 46 and 174 cells).

In CA1, place cells that had place fields in room 4 and 5 showed significantly higher SWR reactivation ratios than those of the other cells in each room (Fig. 4C, bottom; room 1: *n* = 98 cells, –3.24 to 0.60; room 2: *n* = 67 cells, –1.72 to 2.21; room 3: *n* = 73 cells, –0.55 to 3.31; room 4: *n* = 86 cells, 0.65 to 4.50; room 5: *n* = 68 cells, 0.15 to 4.00, 95% confidence intervals of *Z* values defined by Wilcoxon signed-rank test from 10000 bootstrap samples). Furthermore, these ratios were significantly higher than those of place cells that had place fields in room 1 and 2 (Fig. 4D; room 1 vs. 3: *Z* = 3.03, *P* = 0.025; room 1 vs. 4: *Z* = 4.33, *P* = 1.5 × 10^−4^; room 1 vs. 5: *Z* = 3.50, *P* = 0.0046; room 2 vs. 4: *Z* = 2.92, *P* = 0.035; Mann–Whitney U test followed by Bonferroni correction). The same analyses were applied to the CA1 neurons that showed place fields in only one room (Fig. S2), confirming a similar tendency of SWR reactivation ratios to be significantly higher in place cells that had place fields in the latter rooms (Fig. S2A, room 1: *n* = 27 cells, –4.47 to –3.25; room 2: *n* = 10 cells, –2.20 to 1.79; room 3: *n* = 9 cells, –2.33 to 1.13; room 4: *n* = 14 cells, 0.16 to 3.31; room 5: *n* = 5 cells, –1.79 to 2.07, 95% confidence intervals of *Z* values defined by Wilcoxon signed-rank test from 10000 bootstrap samples; Fig. S2B, room 1 vs. 4: *Z* = 4.39, *P* = 1.1 × 10^−4^; room 2 vs. 4: *Z* = 3.10, *P* = 0.020, Mann–Whitney U test followed by Bonferroni correction). Linear regression applied to the CA1 place cells revealed that the coefficients *β* were generally higher as the rooms progressed, except for the highest coefficient *β* in room 4 (Fig. 4F, right; *n* = 174 place cells). These results suggest that post-experience SWR-associated reactivation is stronger in CA1 neurons that encode novelty-related information and latest information, similar to CA3 neurons. Notably, the coefficients *β* from room 1 and 2 were less than 0, confirming that information from initial experiences was negatively reactivated in CA1 during post-experience SWRs.

While linear regression revealed that both CA3 and CA1 neurons generally showed higher coefficients *β* from the novel rooms and the latest room, the magnitudes of their coefficients differed considerably between the two areas (Fig. 4F). To quantify these differences, we applied a multiple linear regression to estimate the effects of novelty and temporal distance on SWR-associated reactivation. In this regression, the first predictor vector was the multiplication of spatial representation patterns (similar to Fig. 4E) and a constant variable composed of binary indices indicating novelty (1 or 0) in each room, and the second predictor vector was the multiplication of spatial representation patterns and a constant variable composed of an exponential decay function over time (Fig. 4G). The regression yielded the coefficients *βnovel* and *βtemp*, representing the effects of novelty and temporal decay, respectively. CA3 place cells showed considerably higher *βnovel* than CA1 place cells (Fig. 4H, left). In contrast, CA1 cells showed a higher *βtemp* than CA3 cells (Fig. 4H, right). Taken together, these results suggest that increases in the post-experience SWR-associated reactivation of CA3 place cells are primarily determined by the novelty of the environment represented by the cells, whereas those of CA1 place cells are primarily determined by the temporal distance from when the cells had place fields.

### Post-experience reactivation of place cells in entire post-rest period

To test the changes in spike patterns during the rest periods regardless of whether they were associated with SWRs, we applied the same analyses to all spike patterns from the entire post-rest period by computing the reactivation ratio as the ratio of the difference in overall spike rates between the entire pre- and post-rest periods to the sum of these rates (Fig. S3A). Similar to SWR-associated reactivation, the reactivation ratios of CA3 place cells that had place fields in at least one room were significantly higher than those that had no place fields (Fig. S3C, *n* = 115 and 48 cells, *Z* = 2.24, *P* = 0.025, Mann–Whitney U test). The CA1 neurons with no place fields or only one field in the five rooms showed significantly lower spike rates during the post-rest period than during the pre-rest period (Fig. S3D, 0 place field: *n* = 86 cells, *Z* = –3.67, *P* = 2.4 × 10^−4^; 1 place field: *n* = 69 cells, *Z* = –2.30, *P* = 0.021; Wilcoxon signed-rank test), while CA1 neurons that had two or more place fields showed no significant differences between the two rest periods (two place fields: *n* = 44 cells, *Z* = –1.09, *P* = 0.28; three place fields: *n* = 33 cells, *Z* = 0.99, *P* = 0.32; four place fields: *n* = 20 cells, *Z* = –0.86, *P* = 0.39; five place fields: *n* = 12 cells, *Z* = 0.24, *P* = 0.81,Wilcoxon signed-rank test). The reactivation ratios of the CA1 neurons with no place fields in any of the five rooms were significantly lower than those of the CA1 place cells with place fields in at least one room (Fig. S3E; *n* = 86 and 178 cells, *Z* = 4.07, *P* = 4.7 × 10^−5^, Mann–Whitney U test). Similarly, CA3 place cells that had place fields in room 2, 4, and 5 showed significantly higher reactivation ratios than the other corresponding cells in each room (Fig. S3F, top; room 1: *n* = 26 cells, –0.83 to 3.01; room 2: *n* = 6 cells, 0.32 to 2.26; room 3: *n* = 6 cells, –1.59 to 2.23; room 4: *n* = 13 cells, 0.39 to 3.07; room 5: *n* = 8 cells, 2.52 to 2.56, 95% confidence intervals of *Z* values defined by Wilcoxon signed-rank test from 10000 bootstrap samples). CA1 place cells with place fields in room 1 showed significantly lower reactivation ratios than the other cells in each room (Fig. S3H, bottom; room 1: *n* = 98 cells, –4.98 to –1.40; room 2: *n* = 70 cells, –3.41 to 0.38; room 3: *n* = 74 cells, –1.78 to 2.18; room 4: *n* = 86 cells, –0.66 to 3.24; room 5: *n* = 68 cells, –1.87 to 2.14, 95% confidence intervals of *Z* values defined by Wilcoxon signed-rank test from 10000 bootstrap samples) and were significantly lower than those of place cells that had place fields in room 3, 4 and 5 (Fig. S3I; room 1 vs 3: *Z* = 3.31, *P* = 0.0094; room 1 vs 4: *Z* = 4.79, *P* = 1.7 × 10^−5^; room 1 vs 5: *Z* = 3.53, *P* = 0.0041; room 2 vs 4: *Z* = 2.93, *P* = 0.033, Mann–Whitney U test followed by Bonferroni correction). Linear regression analysis showed similar tendencies for the coefficient *β* in the individual rooms to those shown in Fig. 4F (Fig. S3K; *n* = 49 CA3 place cells and 178 CA1 place cells). Multiple regression analysis showed that CA3 place cells had higher *βnovel* and *βtemp* values than CA1 place cells (Fig. S3M). These results suggest that the strength of the overall post-experience reactivation of CA3 place cells during the entire post-rest period, regardless of SWR occurrence, is more strongly influenced by both the novelty of the environment represented by the cells and the temporal distance from when the cells had place fields, compared with CA1 neurons.

### Cooperative reactivation of place cell ensembles during post-experience SWRs

Post-rest reactivation patterns at the cell ensemble level were further examined by measuring the correlation of SWR-associated spike patterns between neuron pairs. A vector was constructed from each cell with entries for the number of spikes in each SWR, and the correlation coefficient of the two vectors obtained from simultaneously recorded neurons was computed in the pre- and post-rest periods (Fig. 5A and 5B). The correlations, including cells that generated spikes in fewer than five SWRs, were computed to be 0. For each neuronal pair, changes in spike correlations from the pre- to the post-rest period were quantified by the differences in Fisher’s z-transformed correlations (ΔFisher’s *Z*) (Fig. 5C). The distributions of ΔFisher’s *Z* were separately analyzed for place cell pairs (P-P pairs) defined in each of the five rooms (Fig. 5D and 5F). In CA3, P-P pairs defined in room 4, a novel room, showed significantly higher ΔFisher’s *Z* than the other cell pairs (Fig. 5D; room 1: *n* = 41 and 792 cell pairs, *Z* = 0.59, *P* = 0.55; room 2: *n* = 0 cell pairs (NA); room 3: *n* = 0 cell pairs (NA); room 4: *n* = 12 and 821 cell pairs, *Z* = 4.19, *P* = 2.8 × 10^−5^; room 5: *n* = 4 and 829 cell pairs, *Z* = 0.94, *P* = 0.35, Mann–Whitney U test). Because our datasets had no P-P pairs of CA3 neurons identified in room 2, we defined novelty-encoding P-P pairs by combining the P-P pairs defined in room 4 with a pair of CA3 neurons composed of one cell showing place fields in room 2 and another cell showing place fields in room 4. Overall, CA3 novelty-encoding P-P pairs had significantly higher ΔFisher’s *Z* than the cell pairs encoding the familiar rooms (combined from room 1, 3, and 5) (Fig. 5E; *n* = 12 and 45 cell pairs in the novel rooms and the familiar rooms, respectively, *Z* = 3.36, *P* = 7.7 × 10^−4^, Mann–Whitney U test). These results confirm that correlational spikes of CA3 neurons during post-experience SWRs increase after encoding novel information, consistent with previous reports^22^, but they are not prominently affected by the temporal distance from when the neurons had place fields.

**Figure 5.**
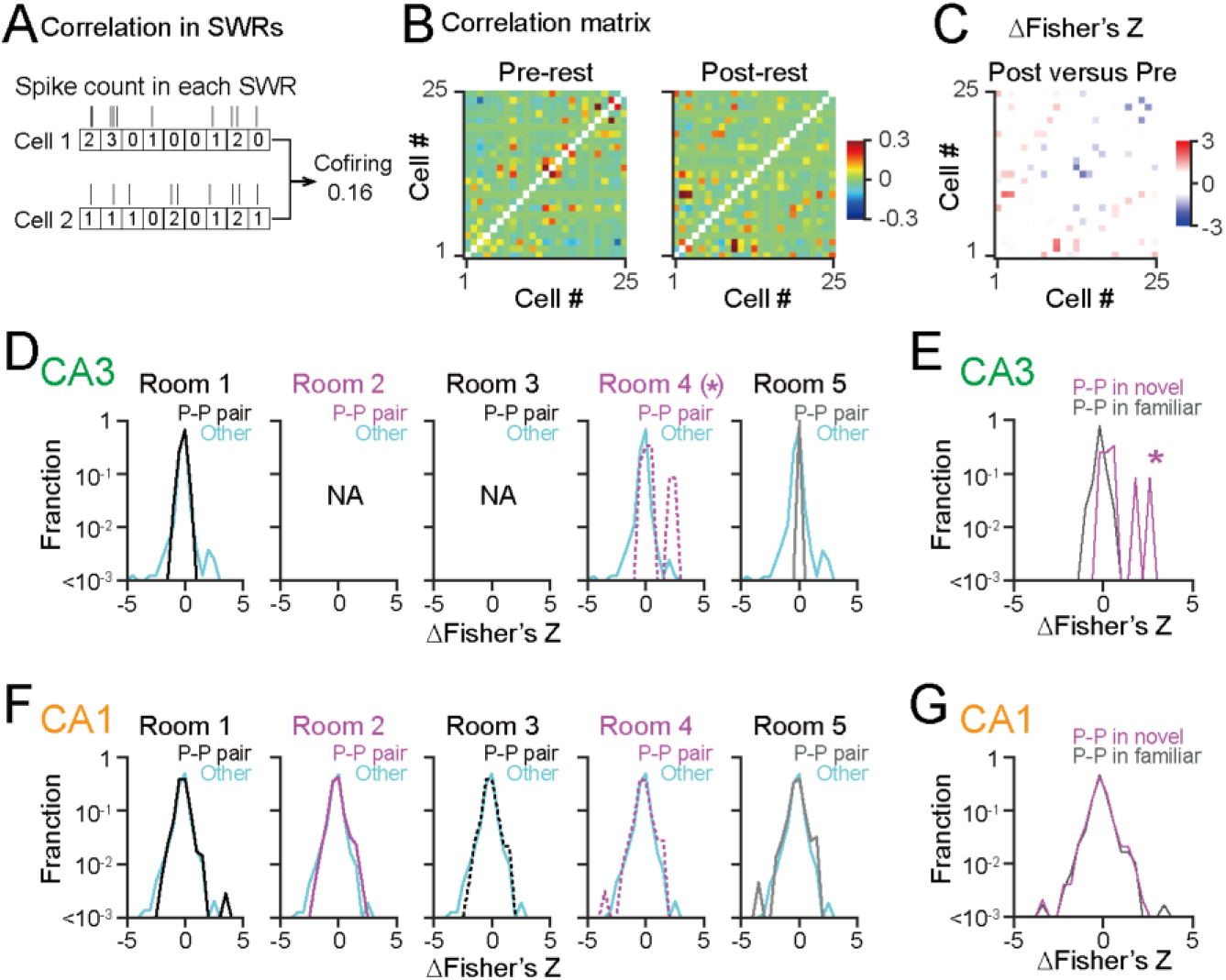
Correlational spikes of place cell pairs during post-experience SWRs. (A) A correlation of spike counts within each SWR was computed for a neuron pair. (B) Correlation matrices for all pairs of neurons computed from the pre- and post-rest periods in a rat. (C) A matrix showing ΔFisher’s *Z* from the two matrices in B. (D) Distributions of ΔFisher’s *Z*, separately created for CA3 neuron pairs that had place fields in each of the five rooms (room 1: *n* = 41 and 792 cell pairs; room 2: *n* = 0 cell pairs (NA); room 3: *n* = 0 cell pairs (NA); room 4: *n* = 12 and 821 cell pairs; room 5: *n* = 4 and 829 cell pairs). Black or magenta lines represent neuron pairs that had place fields in that room (P-P pair) and cyan lines represent the other neuron pairs. \**P* < 0.05, Mann–Whitney U test. (E) Comparison of ΔFisher’s *Z* of CA3 neuron pairs that had place fields in the novel and familiar rooms (*n* = 12 and 45 cell pairs). \**P* < 0.05, Mann–Whitney U test. (F) Same as D but for CA1 neurons (room 1: *n* = 351 and 1452 cell pairs; room 2: *n* = 215 and 1588 cell pairs; room 3: *n* = 230 and 1573 cell pairs; room 4: *n* = 321 and 1482 cell pairs; room 5: *n* = 219 and 1584 cell pairs). (G) Same as E but for CA1 neurons (*n* = 481 and 595 cell pairs).

In CA1, P-P pairs identified from in each of the five rooms showed no significant differences in ΔFisher’s *Z*, compared with the corresponding other cell pairs (Fig. 5F; room 1: *n* = 351 and 1452 cell pairs, *Z* = –0.83, *P* = 0.40; room 2: *n* = 215 and 1588 cell pairs, *Z* = –0.23, *P* = 0.82; room 3: *n* = 230 and 1573 cell pairs, *Z* = –1.12, *P* = 0.26; room 4: *n* = 321 and 1482 cell pairs, *Z* = –0.19, *P* = 0.060; room 5: *n* = 219 and 1584 cell pairs, *Z* = –0.32, *P* = 0.75, Mann–Whitney U test). Similarly, no significant difference in ΔFisher’s *Z* was found between CA1 P-P pairs in the novel and familiar rooms (Fig. 5G; *n* = 481 and 595 cell pairs in the novel and familiar rooms, respectively; *Z* = 0.18, *P* = 0.86, Mann–Whitney U test). These results suggest that changes in SWR-associated correlational spikes in CA1 place cell pairs are not strongly affected by the novelty of the environment or the temporal distance from when the cells had place fields.

## DISCUSSION

We first confirmed that the CA3 place cells showed orthogonalized spatial representations across the five rooms, whereas the CA1 place cells showed overlapping spatial representations across multiple environments ^5,7,8^ (Fig. 6). This spatial encoding of CA1 neurons was generated in cooperation with the other place cell ensemble. During SWRs in subsequent rest periods, the CA3 place cells that showed spatial representations in novel environments were more strongly and cooperatively reactivated than those in familiar environments, in accordance with a previous study ^27^. On the other hand, SWR-associated reactivation of CA1 place cells encoding later environments was more strongly affected by the temporal distance from when the cells had place fields, with little influence from the novelty of the environments, compared with CA3 place cells. No pronounced changes were observed in the spike correlations of CA1 neurons during post-rest SWR-associated reactivation (Fig. 6).

**Figure 6.**
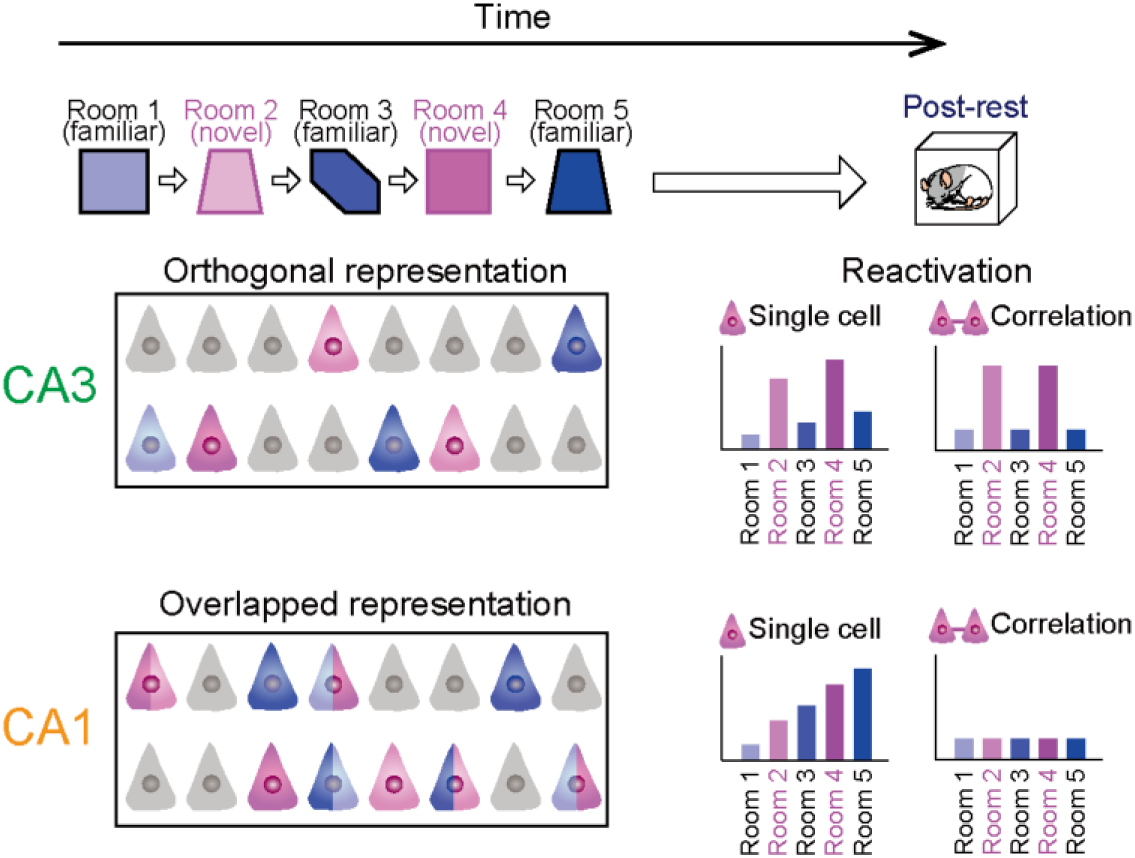
Schematic illustration summarizing the results. (Left) Spatial representations in multiple rooms. Place cells identified from the five rooms are labeled in corresponding colors. The cells that had place fields in multiple rooms are represented by two colors. Non-place cells are colored in gray. (Right) During post-experience SWRs, CA3 place cells in novel environments are more strongly and coordinately reactivated with the other neurons, compared with those in familiar environments. On the other hand, the strength of reactivation of CA1 neurons crucially depends on the temporal distance from when the neurons showed place fields, independent of the other neurons. The novelty of the environments in which neurons showed place fields had weaker effects on reactivation of CA1 neurons, compared with CA3 neurons.

The presence of CA1 place cell ensembles that consistently encode the same places across different environments suggests that these cells form overlapping spatial maps to store multiple spatial memories. This encoding feature can efficiently reduce the sparsity and variability in the spatial representation patterns of neuronal ensembles in multiple environments. Possible mechanisms underlying these ensemble encoding patterns are common synaptic inputs into the CA1 neuronal subpopulation from the CA3 area, with recurrent networks processing orthogonalized spatial information ^5,7,8,34^ and/or the temporo-ammonic pathway from the entorhinal cortex processing multimodal spatial information ^35^. In addition, the interconnections of pyramidal cells and feed-forward and feedback networks that emit common inhibitory inputs within the CA1 circuit may be effective in sustaining specific CA1 place cell ensembles that encode similar information.

Our results suggest that information related to novelty, rather than temporal distance, is prioritized by post-experience SWR-associated neuronal reactivation in the CA3 circuit. This feature potentially embodies effective mechanisms for consolidating novelty-related memories within the hippocampal CA3 circuit ^24,27,36^, the site for SWR generation and initial memory storage. Possible mechanisms underlying novelty-biased information processing by CA3 neurons include NMDA receptor-dependent synaptic plasticity ^37^ and the abundant transmission of neuromodulatory signals from the locus coeruleus ^36,38^. In contrast to CA3, we found that the strength of post-experience neuronal reactivation at the single-neuron level in CA1 was primarily determined by temporal distance, whereas the novelty of the environment had a more modest influence than in CA3. At the same time, we observed no pronounced changes in the spike associations of CA1 place cells during the SWR-associated reactivation. These results demonstrate that CA1 place cells increase their excitability during post-experience SWR-associated reactivation, depending on the temporal distance they encode. However, their reactivated patterns were not strongly coordinated at the cell ensemble level. These observations are different from those observed in CA1 neurons that encode information related to reward ^23^, suggesting that reward-related information serves as a stronger driver of coordinated reactivation in subsequent rest periods than temporal distance-related information. Specifically, CA1 neurons that had no place field or encoded the first environment exhibited decreased reactivation during the post-experience SWRs, whereas those that encoded more recent environments exhibited increased reactivation. These results suggest that distant memory is interfered with or overwritten by recent memory through post-experience CA1 SWR-associated neuronal reactivation. The direction of experience-induced changes in the strength of SWR-associated neuronal reactivation may be counterbalanced throughout the CA1 circuit. The mechanisms underlying elapsed time-dependent reactivation can be attributed to plastic changes or fluctuations in neuronal codes that accumulate over time and experience.

The distinct features of reactivation related to temporal information between CA3 and CA1 neurons share similarities with their spatial encoding characteristics: CA3 neurons can generate highly reproducible spatial encoding over time, whereas CA1 neurons exhibit time-varying spatial encoding, even in the same environments ^9,10^. Taken together, the CA3 circuit ensures both online encoding and offline reactivation of novelty-related information, with modest influence from temporal distance. Conversely, the CA1 circuit is more suitable for both the encoding and reactivation of time-related information, with a weaker influence of novelty.

## LIMITATION OF THE STUDY

In this study, the order of the five rooms, including the familiar and novel environments, was constant for all recordings. When tested in a different order of rooms, the strength of reactivation may vary to some extent, whereas we expect that the overall effects of novelty and room order on subsequent neuronal reactivation may not significantly change. Recent studies have demonstrated that SWR-associated neuronal synchrony involves spike sequences of replays of behavior ^23,39-42^. In this study, owing to the limitations of the spike samples, our spike analysis did not analyze the fine orders of spikes at millisecond timescales.

## ACKNOWLEDGMENTS

This work was supported by KAKENHI (20H03545; 21H05243) from the Japan Society for the Promotion of Science (JSPS), grants (1041630; JP21zf0127004) from the Japan Agency for Medical Research and Development (AMED), grants from the Japan Science and Technology Agency (JST) (JPMJCR21P1; JPMJMS2292) to T. Sasaki; grants from the JST Exploratory Research for Advanced Technology (JPMJER1801), and Institute for AI and Beyond of the University of Tokyo to Y. Ikegaya; and a JSPS Research Fellowship for Young Scientists to Y. Shikano and H. Yagishita.

## AUTHOR CONTRIBUTIONS

T.Y. and T.S. designed the study. Y.S. performed the surgery. Y.S. and H.Y. acquired electrophysiological data and performed spike sorting. T.Y. and T.S. performed the analysis and prepared all figures. Y.I. supervised the project. T.S. wrote the main manuscript text, and all authors reviewed the main manuscript text.

## INCLUSION AND DIVERSITY

We support inclusive, diverse, and equitable conduct of research.

## FIGURE LEGENDS

**Figure S1.**
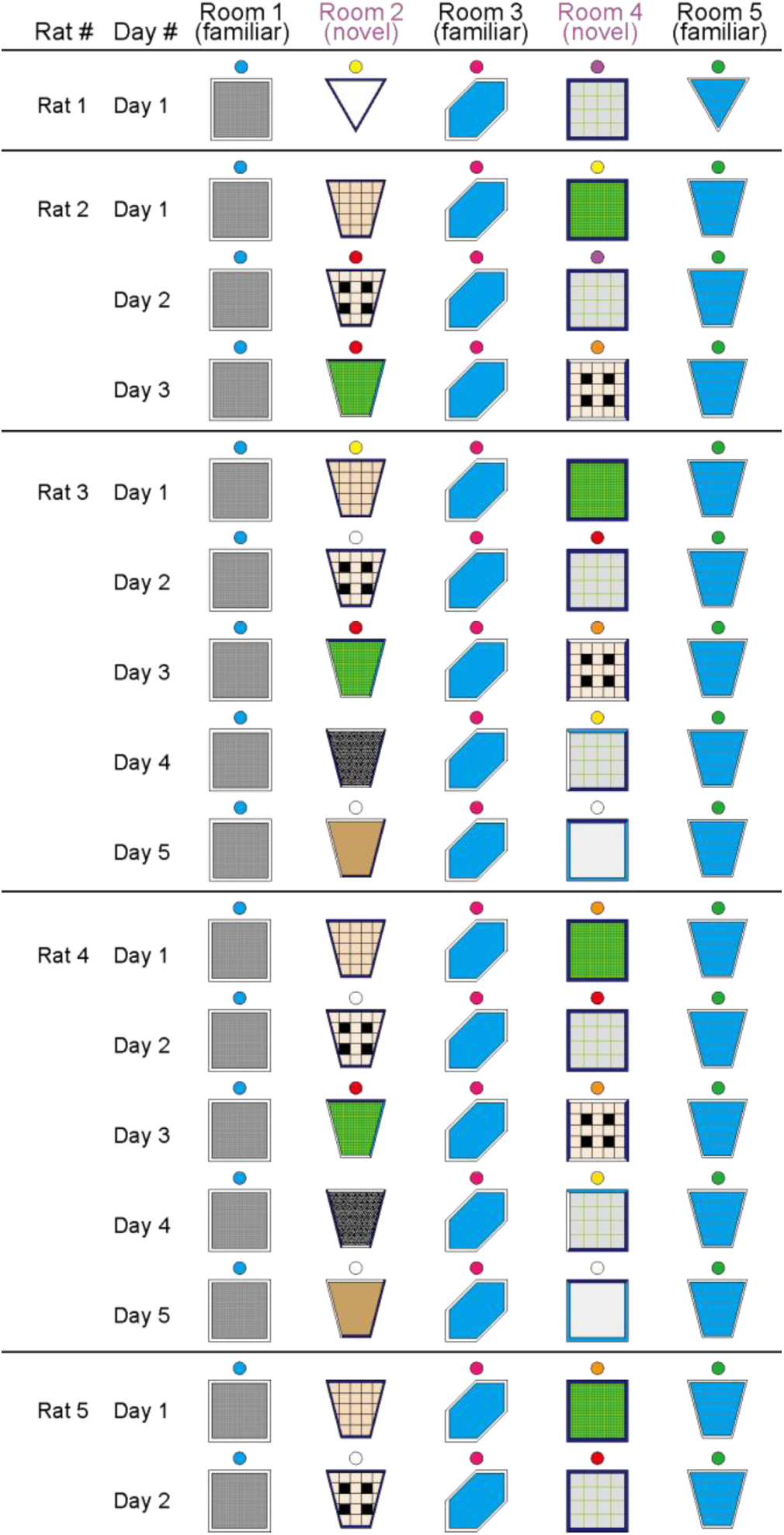
Schematic illustration of recording environments in five rooms. The filled colors and patterns represent the floors of the rooms. The colors of the circles above the rooms represent lights to illuminate the rooms. All rooms were placed in different areas separated by curtains, ensuring that all external cues around the rooms were different. Room 1, 3, and 5 were always similar across all recording days so that the rats were familiar with these rooms. Room 2 and 4 were novel rooms as they had different environments, including floor patterns, lights, and external cues, across recording days.

**Figure S2.**
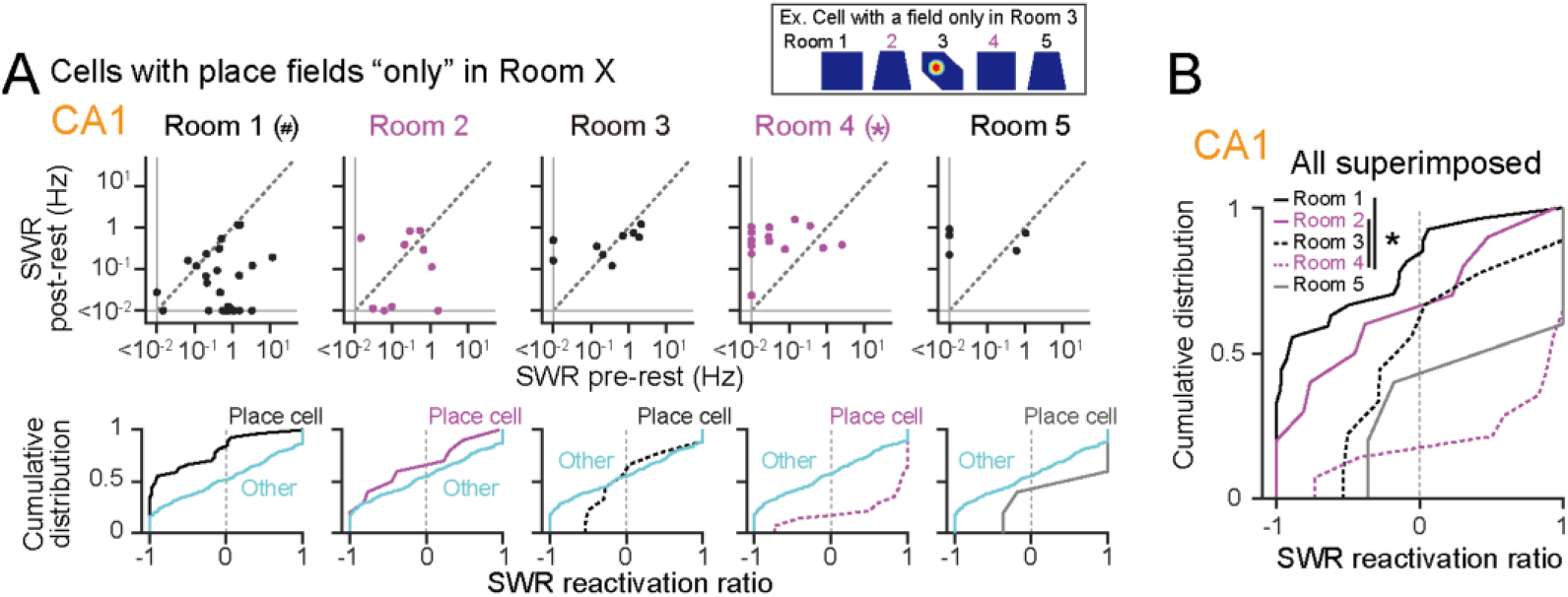
Same as Figure 4C and 4D but for CA1 neurons that showed place fields only in one room. (A) (Top) Scatter plots of SWR-associated spike rates of CA1 neurons during the post-rest period against those during the pre-rest period, which are separately created for neurons that had place fields only in that room (room 1: *n* = 27 cells; room 2: *n* = 10 cells; room 3: *n* = 9 cells; room 4: *n* = 14 cells; room 5: *n* = 5 cells). In this categorization, the neurons that additionally had place fields in the other rooms were classified as the other neurons. The asterisk (*) represents that the lower limit of the 95% confidence interval of *Z* values was more than 0 (post larger), defined by Wilcoxon signed–rank test between pre and post from 10000 bootstrap samples. The hash tag (^#^) represents that the higher limit of the 95% confidence interval of *Z* values was less than 0 (pre larger), defined by Wilcoxon signed-rank test between pre and post from 10000 bootstrap samples. (Bottom) The corresponding cumulative distributions of SWR reactivation ratios. (B) All distributions shown in A are superimposed. \**P* < 0.05, Mann–Whitney U test followed by Bonferroni correction.

**Figure S3.**
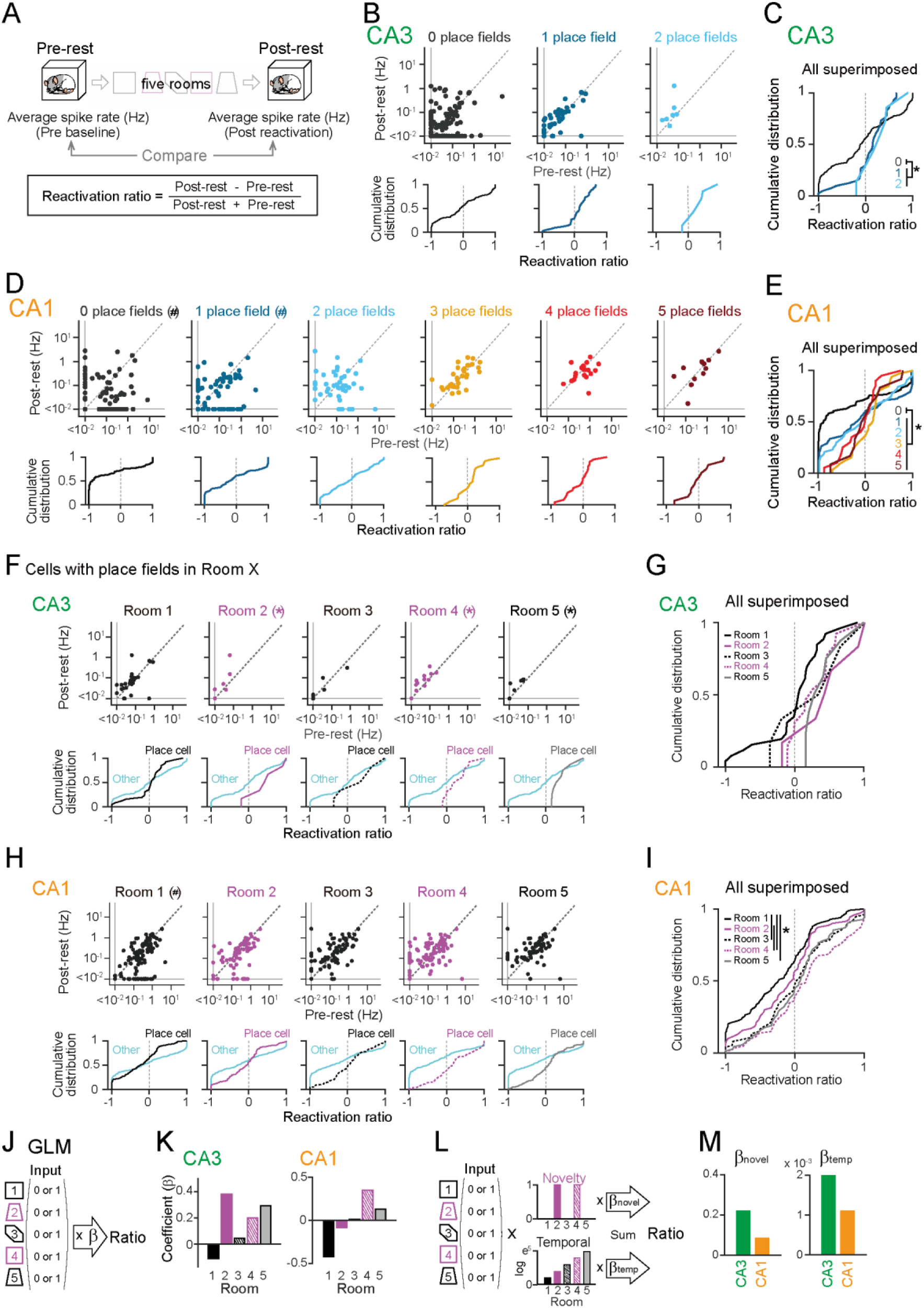
Reactivation of place cells during entire post-rest period. (A) Spike rates during the entire pre- and post-rest periods were compared. A reactivation ratio was computed from each neuron. (B) (Top) Scatter plots of overall spike rates of CA3 neurons during the post-rest period against those during the pre-rest period, separately created for each number of rooms (0: *n* = 105 cells; 1: *n* = 41 cells; 2: *n* = 7 cells). \**P* < 0.05 (post larger), Wilcoxon signed-rank test. (Bottom) Cumulative distributions of reactivation ratios computed from the scatter plots. (C) All distributions shown in B are superimposed. \**P* < 0.05, 0 versus 1–2, Mann–Whitney U test. (D, E) Same as B and C but for CA1 neurons (0: *n* = 86 cells; 1: *n* = 69 cells; 2: *n* = 44 cells; 3: *n* = 33 cells; 4: *n* = 20 cells; 5: *n* = 12 cells). ^#^*P* < 0.05 (pre larger), Wilcoxon signed-rank test. \**P* < 0.05, 0 versus 1–5, Mann–Whitney U test. (F) (Top) Scatter plots of overall spike rates of CA3 neurons during the post-rest period against those during the pre-rest period, which are separately created for neurons that had place fields in each of the five rooms (room 1: *n* = 26 cells; room 2: *n* = 6 cells; room 3: *n* = 6 cells; room 4: *n* = 13 cells; room 5: *n* = 8 cells). The asterisk (*) represents that the lower limit of the 95% confidence interval of *Z* values was more than 0 (post larger), defined by Wilcoxon signed– rank test between pre and post from 10000 bootstrap samples. (Bottom) The corresponding cumulative distributions of reactivation ratios. Black or magenta lines represent neurons that had a place field in that room and cyan lines represent the other neurons that had no place fields. (G) All distributions shown in F are superimposed. (H) Same as F but for CA1 neurons (room 1: *n* = 98 cells; room 2: *n* = 70 cells; room 3: *n* = 74 cells; room 4: *n* = 86 cells; room 5: *n* = 68 cells). The hash tag (^#^) represents that the higher limit of the 95% confidence interval of *Z* values was less than 0 (pre larger), defined by Wilcoxon signed-rank test between pre and post from 10000 bootstrap samples. (I) Same as G but for CA1 neurons. \**P* < 0.05, Mann–Whitney U test followed by Bonferroni correction. (J) The reactivation ratios were linearly regressed against binary-coded spatial representation patterns. (K) Regression coefficients *β* for the individual rooms computed from CA3 and CA1 place cells (*n* = 49 and 178 cells). (L,M) Using multiple linear regression, coefficients *βnovel* and *βtemp* were computed from CA3 and CA1 place cells (*n* = 49 and 178 cells).

**Table S1.** The numbers of simultaneously recorded putative pyramidal cells from the CA1 and CA3 regions.

| Rat ID | Experiment | N of putative excitatory cells |
| --- | --- | --- |
| Rat 1 | CA1, Day 1 | 39 |
| Rat 2 | CA1, Day 1 | 12 |
|  | CA1, Day 2 | 21 |
|  | CA3, Day 1 | 5 |
| Rat 3 | CA1, Day 1 | 24 |
|  | CA1, Day 2 | 29 |
|  | CA1, Day 3 | 31 |
|  | CA3, Day 1 | 10 |
|  | CA3, Day 2 | 28 |
| Rat 4 | CA1, Day 1 | 26 |
|  | CA1, Day 2 | 40 |
|  | CA3, Day 2 | 34 |
|  | CA3, Day 2 | 35 |
|  | CA3, Day 3 | 54 |
| Rat 5 | CA1, Day 1 | 5 |
|  | CA1, Day 2 | 37 |

## Methods

### Resource availability

#### Lead contact

Further information and requests for resources and reagents should be directed to and will be fulfilled by the lead contact, Takuya Sasaki.

### Materials availability

This study did not generate any unique reagents. The CAD files for creating the tetrode assembly by 3D printers are available from the lead contact upon request.

### Data and code availability

• The original data and codes are available from the lead contact upon request.

• Any additional information required to reanalyze the data reported in this paper is available from the lead contact upon request.

Experimental model and study participant details

## Experimental model and subject details

All experiments were performed with the approval of the experimental animal ethics committee at the University of Tokyo (approval number: P29-7) and according to the NIH guidelines for the care and use of animals.

All Long Evans male rats were purchased from SLC (Shizuoka, Japan) and 3 male Long Evans rats (3–6 months old) with a preoperative weight of 400–500 g were used in this study. The animals were housed individually and maintained on a 12-h light/12-h dark schedule with lights off at 7:00 AM. Following at least 1 week of adaptation to the laboratory, the rats were reduced to 85% of their *ad libitum* weight through limited daily feeding. Water was readily available.

## Method details

### Behavioral habituation to familiar fields before surgery

All behavioral experiments were conducted in the dark. Before surgery, rats were trained to perform random foraging in three different rooms (room 1, 3, and 5). Each room was positioned at a different location and orientation, all separated by black curtains with different global cues (e.g., posters and walls). Room 1 was a 1-meter square with a white floor and blue lighting. Room 3 had a hexagonal shape with two 90-degree angles and four 135-degree angles, situated within a 1-meter square area with a blue floor and pink lighting. Room 5 was in the form of a trapezoid with an upper base of 0.5 meters, a lower base of 1 meter, and a height of 1 meter, featuring a blue floor and green lighting. All rooms had a wall height of 20 cm and were elevated 50 cm above the floor.

On the training day, the rats first rested in a rest box (30 × 30 cm) outside the rooms for 30 min and then sequentially experienced the three rooms (room 1, 3, and 5). In each session, the rats were placed in a room and habituated to the environment by allowing them to freely forage for randomly placed chocolate milk samples for 10 min. Between sessions, they returned to the same rest box for 5 min. After three sessions, the rats rested in the same box for 30 min. This training was repeated daily until the rats consumed the reward at least 20 times in a 10-min session. To achieve this criterion, the training lasted for at least four days.

### Surgical procedures

The rat was anesthetized with isoflurane gas (0.5–2.5%), and a 2-cm midline incision was made in the area between the eyes and the cerebellum. A craniotomy with a diameter of 1.5 mm was created above the right dorsal hippocampus (3.5 mm posterior and 3.3 mm lateral to the bregma) using a high-speed drill, and the dura was surgically removed. Two stainless steel screws were implanted in the bone above the prefrontal cortex to serve as ground electrodes. An electrode assembly consisting of 13–15 independently movable tetrodes was stereotaxically implanted above the craniotomy ^15,43,44^. The tips of the tetrode bundles were lowered to the cortical surface, and the tetrodes were inserted 1.0 mm into the brain at the end of surgery. The tetrodes were constructed from 17-μm-wide polyimide-coated platinum-iridium (90/10%) wire (California Fine Wire California Fine Wire Co., Grover Beach, CA) and plated with platinum to reduce their electrode impedances to 150–300 kΩ at 1 kHz. All recording devices were secured to the skull using stainless-steel screws and dental cement. Following surgery, each rat was housed individually in a transparent Plexiglass with free access to water and food for at least 5 days and was then food-deprived until they reached 85% of their previous body weight.

### Adjusting electrode depth

The rat was connected to the recording equipment via Cereplex M (Blackrock), a digitally programmable amplifier, close to the rat’s head. The output of the headstage was conducted via a lightweight multiwire tether and a commutator to the Cereplex Direct recording system (Blackrock), a data acquisition system. Electrode turning was performed while the rat was resting in a rest box. Over a period of at least 2 weeks after surgery, tetrode tips were advanced slowly 25–100 μm per day for 14–21 days until spiking cells were encountered in the CA1 layer of the hippocampus, which was identified on the basis of local field potential (LFP) signals and single-unit spike patterns. Once the tetrodes were adjacent to the CA1 cell layer, as indicated by the presence of low-amplitude multiunit activity, tetrodes were settled for stable recordings over a period of several days. After recordings from CA1 cells, the tetrodes were further advanced slowly 25–100 μm per day for 14–21 days until spiking cells were encountered in the CA3 layer of the hippocampus, which was identified on the basis of LFP signals and single-unit spike patterns. Once the tetrodes were adjacent to the CA3 cell layer, recordings were obtained over a period of several days.

### Recoding paradigm

After surgery, behavioral habituation resumed at least 5 days before the recording day. Electrophysiological recordings were conducted after the rats consumed the reward at least 20 times within the 10-min session for at least three consecutive days. On the recording day, the rats initially rested in the box for 30–60 min (pre-rest) and then sequentially explored five rooms (room 1, 2, 3, 4, and 5). Finally, the rats rested in a rest box for 30–60 min (post-rest). Similar to the training, the rats freely foraged in each chamber for 10 min and rested for 5 min between sessions, during which the floor of the field was cleaned with water and 70% ethanol. Room 2 and 4 were novel, with trapezoidal and square shapes, respectively (except for rat 1). All rooms were positioned in different locations and orientations and were separated by black curtains with different global cues (e.g., room poster and walls). Every day, these novel rooms featured distinct flooring materials such as polyurethane, plastic, corrugated board, or artificial turf, along with cues provided by green tape, black tape, a black star, or red and yellow squares. The walls were either blue or white, and the lighting was in various colors, including yellow, orange, purple, red, and white. The rooms had wall heights of 30 or 60 cm and were elevated 50 cm above the floor.

### Electrophysiological data collection

Electrophysiological data collection commenced after stable well-separated unit activity was identified in the hippocampus and the rat reached the criterion performance. LFP recordings were sampled at 2 kHz and lowpass filtered at 500 Hz. Unit activity was amplified and highpass filtered at 750 Hz. Spike waveforms above a trigger threshold (50 μV) were timestamped and recorded at 30 kHz for 1.6 ms. To monitor the rat’s moment-to-moment position, an infrared light reflective tape was attached to the microdrive, and the LED signal position was tracked at 25 Hz using an infrared camera located on the ceiling and sampled by a laptop computer. All the recording equipment placed between the recording room and the room entrance was wheeled across rooms without stopping the recordings to ensure identical recording conditions (e.g. sampling, filtering and, amplification). Recordings were conducted for at least 2 days.

### Histological analysis to confirm electrode placement

After the experiments, the rats received an overdose of urethane and were intracardially perfused with 4% paraformaldehyde (PFA) in phosphate buffered saline and decapitated. To aid in the reconstruction of the electrode tracks, the electrodes were not withdrawn from the brains until more than 3–4 hours after perfusion. After dissection, the brains were fixed overnight in 4% PFA and then equilibrated with 30% sucrose in PBS. Frozen coronal slices (50 μm) were cut using a microtome, and serial sections were mounted. The slices were rinsed in water, counterstained with cresyl violet, and coverslipped with hydrophobic mounting medium. The positions of all the tetrodes were confirmed by identifying the corresponding electrode tracks in histological tissue with an optical microscope.

## Quantification and statistical analysis

### Spike sorting

Spike sorting was performed offline using the graphical cluster-cutting software MClust. Rest recordings before and after the behavioral paradigms were included in the analysis to assure recording stability throughout the experiment and to identify hippocampal cells that were silent during behavior. Clustering was performed manually in 2D projections of the multidimensional parameter space (i.e., comparisons between waveform amplitudes, the peak-to-trough amplitude differences, and waveform energies, each measured on the four channels of each tetrode). The cluster quality was measured by computing the Lratio and isolation distance. A cluster was considered as a cell when the Lratio was less than 0.38 (average Lratio was 0.105 ± 0.006 in 430 isolated cells). In the auto-correlation histograms, cells with no clear refractory period (<3 ms) were excluded from analyses. Refractory periods of spikes were considered to increase confidence in the successful isolation of cells. In addition, in the cross-correlation histograms, putative cell pairs with a symmetrical gap around the center bins were considered to arise from the same cell and were merged.

### Analysis of spatial spike patterns

The following analyses were applied for each cell. For quantifying spatial selectivity, a spatial firing-rate distribution was constructed by dividing the sum of the total number of spikes in each location bin (7 cm × 7 cm) by the amount of time that the animal spent in that bin. Data were smoothed with a Gaussian kernel filter with a standard deviation of 1.5 pixel (11 cm), constructing a tuning curve map. A spatial information density was computed by the following formula:

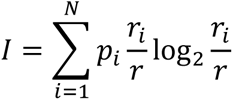

where *I* is the spatial information density measured in bits per spike, *i* is the index of the pixels of the place field, *pi* is the probability of the animal being at location *i*, *ri* is the average firing rate of the cell when the animal is at location *i*, and *r* is the total average firing rate. A cell was classified as a place cell if the following criteria were met: (1) the spatial information was more than 0, (2) the spatial information was more than 1% of the top of those computed from 100 randomized data in which spike times were shifted along the recording period by a random amount within the duration of the recording session, (3) the average firing rate in the room was more than 0.05 Hz, and (4) the maximum firing rate among all bins was more than 1 Hz.

### Detection of synchronous events and SWRs

The electrode including the largest number of putative pyramidal cells identified in the spike sorting process was used for SWR detection. LFP signals during rest periods were bandpass filtered at 150–250 Hz, and the root mean-square was calculated with a bin size of 10 ms. SWR events were detected if the power exceeded a threshold for at least 20 ms. The threshold for SWR detection was set to 3 standard deviations (SDs) above the mean of all envelopes computed from the rest periods. SWR events that occurred within 100 ms from the former SWR events were excluded. The onset of SWRs was marked at the point when the root mean-square first exceeded 3 SDs above the mean. SWR-associated spike rates were computed from –50 to +100 ms relative to SWR onsets. For each cell, a SWR reactivation ratio was computed as the ratio of the difference in its SWR-associated spike rates between the pre-rest and post-rest periods to the sum of these rates. The ratios ranged between –1 and +1.

### Linear regression of SWR reactivation ratios from spatial representation patterns

For the *l*-th place cell, a (5+1)-dimensional predictor vector *X*_*l*_ consists of five-dimensional entries, each representing a binary index indicating the presence (1) or absence (0) of its place fields in each room. The (5+1)-th entry was set to +1 as a constant term. The dependent variable *Y*_*l*_ was set as the SWR reactivation ratio. A linear relationship between a series of predictor vectors, *X* = (*X*_1_, *X*_2_, *X*_3_,···, *X*_*L*_)^*T*^, and a series of dependent variables, *Y* = (*Y*_1_, *Y*_2_, *Y*_3_,···, *Y*_*L*_)^*T*^, where *L* denotes the total number of place cells, was estimated by a linear regression analysis in which the best (5+1)-dimensional weighted coefficients *β* were mathematically computed so that the weighted linear sum *Y’* (= *X·β*) was fitted against *Y* as follows:

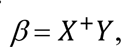

where *X*^+^ = (*X*^*T*^*X*)^−1^*X*^*T*^.

The effects of novelty and temporal distance on SWR-associated reactivation were estimated using multiple linear regressions. For the *l*-th place cell, two predictor vectors were created. The first is a (5+1)-dimensional predictor vector representing novelty *X*_*novel*_*l*_ which is the product of *X*_*l*_ and a constant vector *Novel* = (0, 1, 0, 1, 0, 1), composed of five-dimensional entries, each representing the binary index indicating the novel (1) or familiar (0) rooms. The (5+1)-th entry was set to +1 as a constant term. The second is a (5+1)- dimensional predictor vector representing the decay effects of the temporal distance *X*_*temp*_*l*_ which is the product of *X*_*l*_ and a constant vector *Temp* = (*e*, *e^2^*, *e^3^*, *e^4^*, *e^5^*, 1), composed of five-dimensional entries of an exponential decay function over time, assuming that the effects of reactivation undergo exponential decay. The (5+1)-th entry was set to +1 as a constant term. A linear relationship between two predictor vectors, *X*_*novel* and *X*_*temp*, and the dependent variables *Y* was estimated by a multiple linear regression analysis in which the best (5+1)-dimensional weighted coefficients *βnovel* and *βtemp* were mathematically computed so that the weighted linear sum *Y’* (= *X*_*novel·βnovel* + *X*_*temp·βtemp*) was fitted against *Y*.

### Spike correlations in a cell pair

To measure the degree to which a given cell pair exhibited correlational spikes during SWRs, the numbers of spikes during each SWR was computed from each of the two cells, creating N-dimensional vectors x and y, where N is the total number of SWRs. Pearson correlation coefficients were computed between the two vectors as follows:

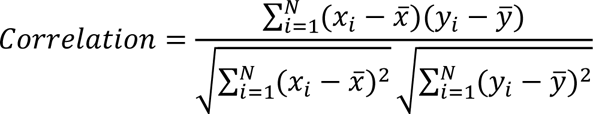

Correlations including cells that generated spikes in less than five SWRs were computed as 0. To evaluate whether a neuronal pair showed a significant difference in correlations from the pre-rest period to the post-rest period (*correlationpre* to *correlationpost*), a difference in Fisher’s z-transformed value was computed as follows:

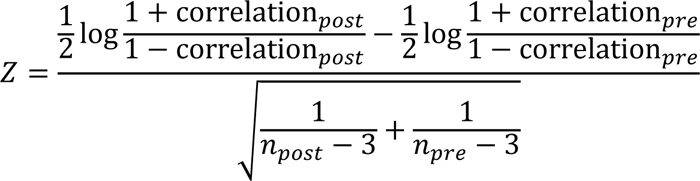

where *npre* and *npost* were the numbers of SWRs computed for *correlationpre* and *correlationpost*, respectively.

### Statistical analysis

All data were analyzed using Python and MATLAB software. Two-sample data were compared using Student’s *t*-test, Mann–Whitney U test, and Wilcoxon signed-rank test. Multiple group comparisons were performed using post-hoc Bonferroni correction. The probabilities of observing place cells were compared using the chi-squared test. The null hypothesis was rejected at *P* < 0.05 level.

When the sample size was too small for statistical tests, 95% confidence intervals of *Z* values were defined using the Wilcoxon signed-rank test from bootstrap samples. The results were considered significant if the lower or higher limit of the 95% confidence interval of *Z* values was more than 0 or less than 0, respectively.

